# Large-scale simulations of biological cell sorting driven by differential adhesion follow diffusion-limited domain coalescence regime

**DOI:** 10.1101/2020.12.15.422842

**Authors:** Marc Durand

## Abstract

Cell sorting, whereby a heterogeneous cell mixture segregates and forms distinct homogeneous tissues, is one of the main collective cell behaviors at work during development. Although differences in interfacial energies are recognized to be a possible driving source for cell sorting, no clear consensus has emerged on the kinetic law of cell sorting driven by differential adhesion. Using a modified Cellular Potts Model algorithm that allows for efficient simulations while preserving the connectivity of cells, we numerically explore cell-sorting dynamics over unprecedentedly large scales in space and time. For a binary mixture of cells surrounded by a medium, increase of domain size follows a power-law with exponent *n* = 1/4 independently of the mixture ratio, revealing that the kinetics is dominated by the diffusion and coalescence of rounded domains. We compare these results with recent numerical and experimental studies on cell sorting, and discuss the importance of boundary conditions, space dimension, initial cluster geometry, and finite size effects on the observed scaling.

**Author summary:** Cell sorting describes the spontaneous segregation of identical cells in biological tissues. This phenomenon is observed during development or organ regeneration in a variety of biological systems. Minimization of the total surface energy of a tissue, in which adhesion strengh between homotypic and heterotypic cells are different, is one of the mechanisms that explain cell sorting. This mechanism is then similar to the one that drives demixing of two immiscible fluids. Because of the high sensibility of this process to finite-size and finite-time effects, no clear consensus has emerged on the scaling law of cell sorting driven by differential adhesion. Using an efficient numerical code, we were able to investigate this scaling law on very large binary mixtures of cells. We show that on long times, cell sorting obeys a universal power law, which is independent of the mixture ratio.

## Introduction

Collective cell behaviors are involved in many morphogenetic events, such as embryo development, organ regeneration, wound healing, and the progression of metastatic cancer [1]. One of the simplest and best studied examples of such behavior is the spontaneous separation of two randomly mixed cell populations, in a process called cell sorting [2]. In the 1960s, Steinberg proposed the differential adhesion hypothesis (DAH) as a mechanism to explain cell sorting [3]. The DAH postulates that the surface tension of tissue arises from cell adhesion, and that cell configurations are determined by minimization of the surface energy of cell aggregates in a manner similar to phase separation processes, although surface energy minimization is an active process for cell sorting: cellular activity is necessary to untrap from the many local minima of the rough energy landscape of a cellular system. Since Steinberg’s hypothesis statement, several numerical studies have proven that the DAH could reproduce cell sorting phenomena similar to those observed experimentally [4–8]. However, no clear consensus has emerged from these studies on the kinetics law of DAH-driven cell sorting : in their seminal papers [4, 5], Graner and Glazier found – on a somewhat smaller system – that the *boundary length*, defined as the number of mismatched cell contacts, decays logarithmically. More recently, Nakajima *et al.* [9] reported a power-law decay, with an exponent which depends on the proportion of the two cell types: from *n* = 1/3 for even mixture to *n* = 1/4 for uneven mixture. Cochet-Escartin *et al.* [8] also reported power-law decays, but with significantly higher exponents: *n* ∈ [0.49, 0.74] from experiments, while *n* ∈ [0.55, 0.59] from simulations.

Meanwhile, alternative mechanisms have been proposed in recent years to explain cell sorting phenomena – such as differences in cell motility that are either intrinsic [10] or dependent on the cells’ local environment [11,12] – leading to different scaling behaviours. In this context, more extensive simulations of DAH-driven cell sorting are needed to find out which type of kinetics law actually governs cell sorting.In this paper, we present extensive DAH-driven cell sorting simulations for two-dimensional binary mixtures of cells surrounded by medium, using a recently modified Cellular Potts Model (CPM) algorithm that allows for simulations on very large scales in space and time. On long times, reported kinetics obeys algebraic scaling law with exponent *n* = 1/4 independently of the mixture ratio. We discuss the importance of boundary conditions, initial cluster geometry, space dimension, and finite size effects on the observed scaling.

## Methods

### Cellular Potts Model

Cellular Potts model (CPM), also called the Glazier–Graner–Hogeweg (GGH) model, is one of the most accepted models of a multicellular system. It was intially created to simulate the behavior of single cells during DAH-driven cell sorting [4, 5]. Since then, this model has been extended to reproduce many other collective behaviors of cells [13–16]. Cellular activity is modeled with an effective temperature *T*, and the update algorithm is based on Metropolis-like dynamics [6]. The main justification for it is that average time evolution of a cell configuration then obeys the overdamped force-velocity relation ***F*** = *μ**v***, where ***v*** is the cell velocity, *μ* is an effective mobility, and 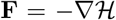 is the acting force [17]. The CPM is a lattice based modeling technique: each cell is a subset of lattice sites sharing the same cell ID *σ*. A cell type *τ* (*σ*) is also defined for each cell, that in our case can have three distinct values corresponding to endoderm, ectoderm and medium, respectively. Following [4, 5], the CPM Hamiltonian 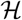 we use reads:

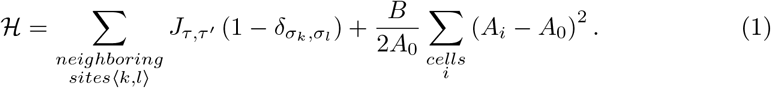

The first sum in Eq. (1) is carried over neighbouring sites 〈*k, l*〉 and represents the boundary energy: each pair of neighbours having unmatching indices determines a boundary and contributes to the boundary energy. Here, *σ_k_* and *σ_l_* are the site values of site *k* and *l*, respectively. *τ* and *τ′* are abbreviations for *τ* (*σ_k_*) and *τ* (*σ_l_*). *J_τ,τ′_* (= *J_τ′_,τ*) is the energy per unit contact length between cell types *τ′* and *τ*. The second sum in Eq. (1) represents the compressive energy of the cells. *B* is the effective bulk modulus of a cell, which captures its out-of-plane elongation [18, 19], *A_i_* is the area of cell *i*, and *A*0 the *nominal area*.

Eq. 1 is the minimal Hamiltonian whose minimization induces cell sorting phenomena. However, various additional terms can be added to account for real cell behaviors, or application of external fields [13, 20–22].

CPM presents a number of advantages over competing numerical models for multicellular systems, such as vertex- or voronoi-based models [23]: interfaces can have arbitrary shapes, T1 topological rearrangements or neighbor exchanges, which are crucial for dynamics in such systems, are naturally included,and free boundaries with the medium are handled as simply as boundaries between adjacent cells. However, a notorious weakness of CPM – when used with the standard updating rule – is that it does not guarantee connectivity of cells: as temperature increases, small fragments detach from cells, biasing the cell perimeters and cell centers of mass. Ideally, CPM cell sorting simulations should be run in a temperature range such that interface fluctuations are high enough for generating T1 neighbors swapping, but low enough for avoiding cell fragmentation. However, such a temperature range is almost nonexistent: the thermal energy required to generate a T1 in a hexagonal pattern (honeycomb) interface fluctuations is estimated to a fraction of the typical surface energy of one side: 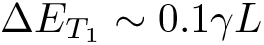 [24]. In comparison, the minimal energy to create a fragment with size *l* is ~ *γl* [25]. Then, fragments of size up to *l* ≃ 0.1*L* show up before first T1 rearrangements occur. To prevent the visual appearance of fragments, quenching process is usually performed before snapshots. However, the creation of fragments, inefficient for the cell sorting process, artificially slows down simulations [25].

### New algorithm that preserves cell connectivity

In a recent work [25], we proposed a new CPM algorithm that forbids fragmentation and improves substantially computational efficiency for a same running temperature: the time wasted in checking cell connectivity is more than offset by the time spared in fragments moves, evidencing that fragmentation occurs even at low temperature. Moreover, efficiency can be further enhanced by working at temperature range that are unattainable with the standard algorithm, due to the proliferation of fragments. Because the dynamics of our algorithm also belongs to the Metropolis-like class, we expect to observe the same kinetics than with the standard algorithm, *i.e.* all things being equal, the real time-correspondence of one MCS for both algorithms should be proportional to each other.

Here we use this efficient algorithm to study DAH-driven cell sorting kinetics on unprecedently large systems. We performed simulations for binary mixtures of *N* cells, with *N* ranging from 200 to 320, 000. Moreover, in order to smooth out completely the anisotropy of the underlying square lattice, we used a 4th order neighborhood (composed of 20 lattice sites) in the evaluation of the Hamiltonian, whereas previous studies used a 2nd order neighborhood, composed of 8 lattice sites only [4, 5, 9]. Consistently, we also used a larger cell size (the target area was set to *A*_0_ = 100 pixels), to avoid interactions between opposite cell sides. Values of the other parameters are: *T* = 50, *B* = 200, *J_BB_* = *J_Y Y_* = 8, *J_Y B_* = 14, *J_Y m_* = 10, *J_Bm_* = 22, where *m* stands for the *medium*, while *B* and *Y* stand for *Blue* and *Yellow*, the colors of the endodermal and ectodermal cells in our graphical outputs, respectively.

## Results

Figure 1 shows different snapshots of the simulation for an even (50:50) mixture of endodermal (blue) and ectodermal (yellow) cells. Initially, the two types of cells were distributed randomly in a rounded cluster (Fig. 1(a)). Subsequently, cells of the same type began to aggregate and form clusters (Fig. 1(b)), and the sizes of the clusters grew with time (Fig. 1(c)).

**Fig 1.**
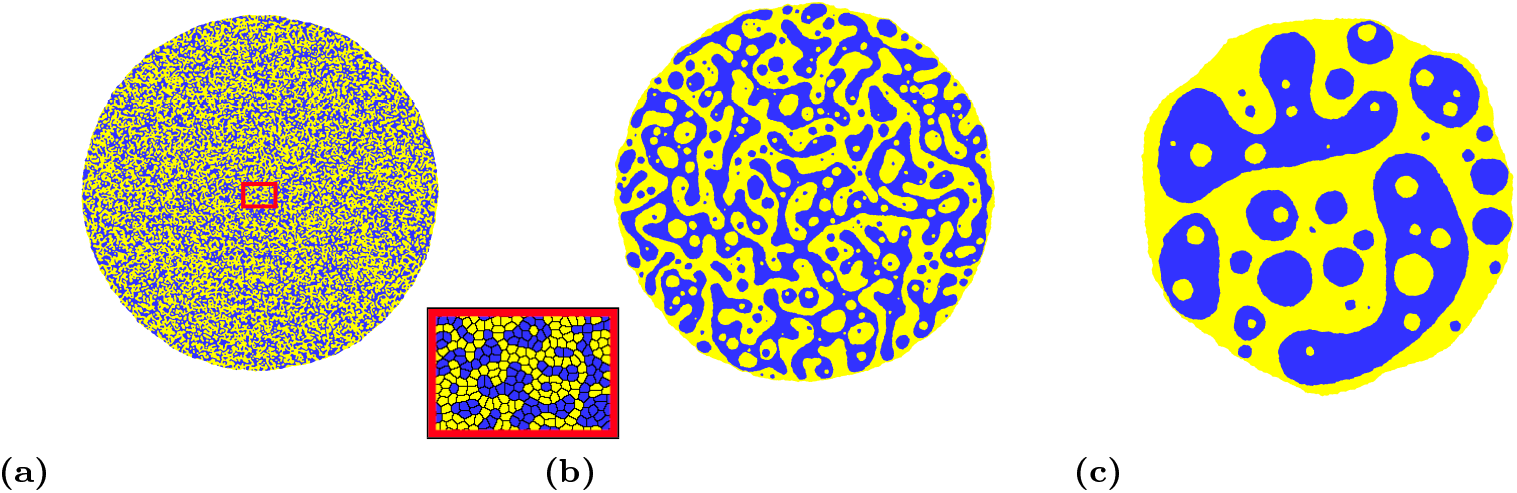
Time evolution of a cluster of 45×10^3^ cells with 50%-50% proportion of endodermal (in blue) and ectodermal (in yellow) cells: (a): t= 10^3^ MCS, (b): t= 40 × 10^3^ MCS, (c): t =2 × 10^6^ MCS. Close-up: zoom on the aggregate area indicated by a red rectangle. Nominal area of each cell is 100 pixels^2^

Choice of running temperature has a strong influence on the cell sorting timescale: fluctuations must be large enough to jump from a local minimum of energy to a lower one, but not too large to avoid jumping to a higher one, so there is likely an optimal temperature for which cell sorting timescale is minimal. Most importantly, there is a full temperature range for which the *kinetics* of cell sorting is preserved. Indeed, the net rate of transition from configuration *i* to configuration *j* obtained with Metropolis-like algorithm is 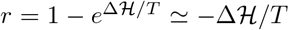 for small relative change of energy 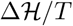. In this regime, a change in temperature uniformly scales the rate of transition between configurations, leaving the kinetics of cell sorting unchanged.

Following previous studies [4, 5, 26], we quantify the kinetics of cell sorting by tracking the evolution of the *total boundary length* Γ(*t*), defined as the number of mismatched cell contacts. Figure 2 shows the evolution of Γ(*t*) for even (50:50) mixtures of various sizes. We see that before complete cell sorting, all the curves follow the same algebraic variation: Γ(*t*) ~ *t^−n^* with *n* ≃ 1/4. Unexpectedly, our exponent value is different from the one reported by Nakajima and Ishihara [9] for a same mixture ratio (*n* = 1/3), but is identical to the value they obtain for uneven mixture ratios.

**Fig 2.**
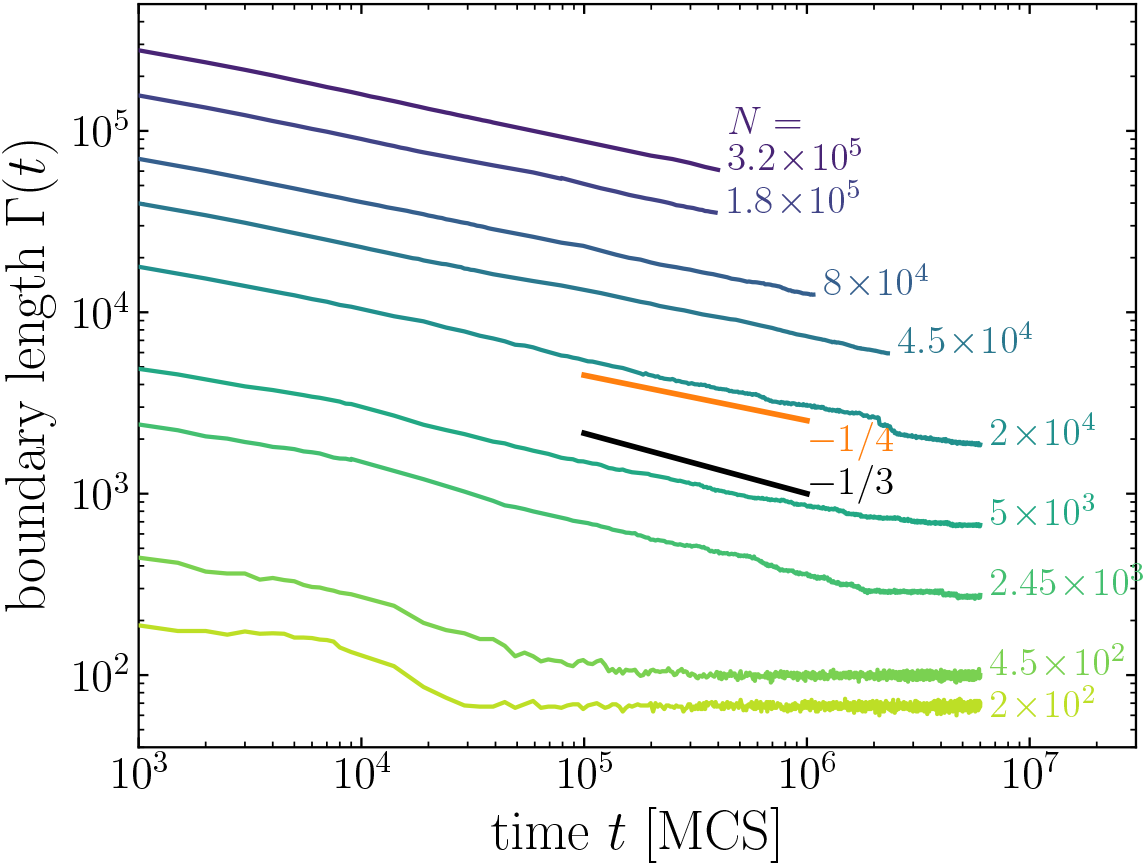
Time evolution of the boundary length Γ(*t*) for even mixtures of *N* endodermal and ectodermal cells, with *N* ranging from 200 to 320, 000 (log-log scale). Power-law exponent is closer to 1/4 than 1/3.

We then study how the mixture ratio affects the power-law exponent in our simulations. Figure 3 shows the typical time evolution of cell sorting for a 20:80 mixture. As for the 50:50 mixture, cells of the same type aggregate into clusters whose size grows with time. We note however that at same simulation time, clusters are much smaller and rounder than for the 50:50 mixture. Fig. 4 compares the time evolution of boundary length for a 50:50 and a 20:80 mixture ratio. Both curves rapidly follow the same power law with exponent *n* = 1/4.

**Fig 3.**
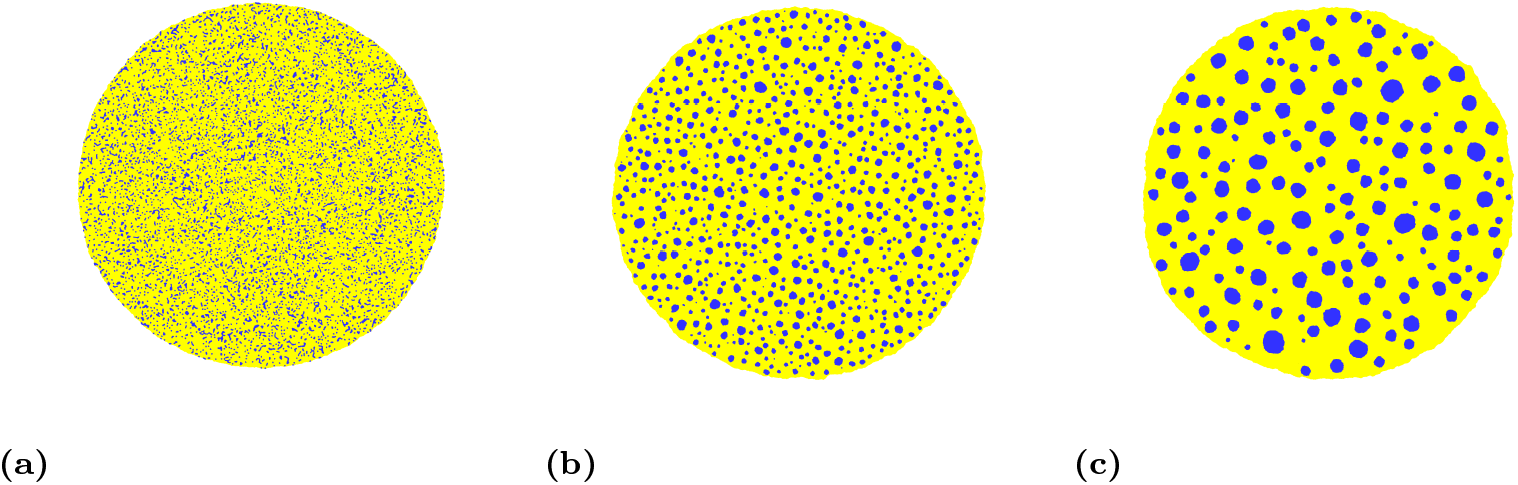
Time evolution of a cluster of 45×10^3^ cells with 20%-80% proportion of endodermal (in blue) and ectodermal (in yellow) cells: (a): t= 10^3^ MCS, (b): t=40 × 10^3^ MCS, (c): t =2 × 10^6^ MCS.

**Fig 4.**
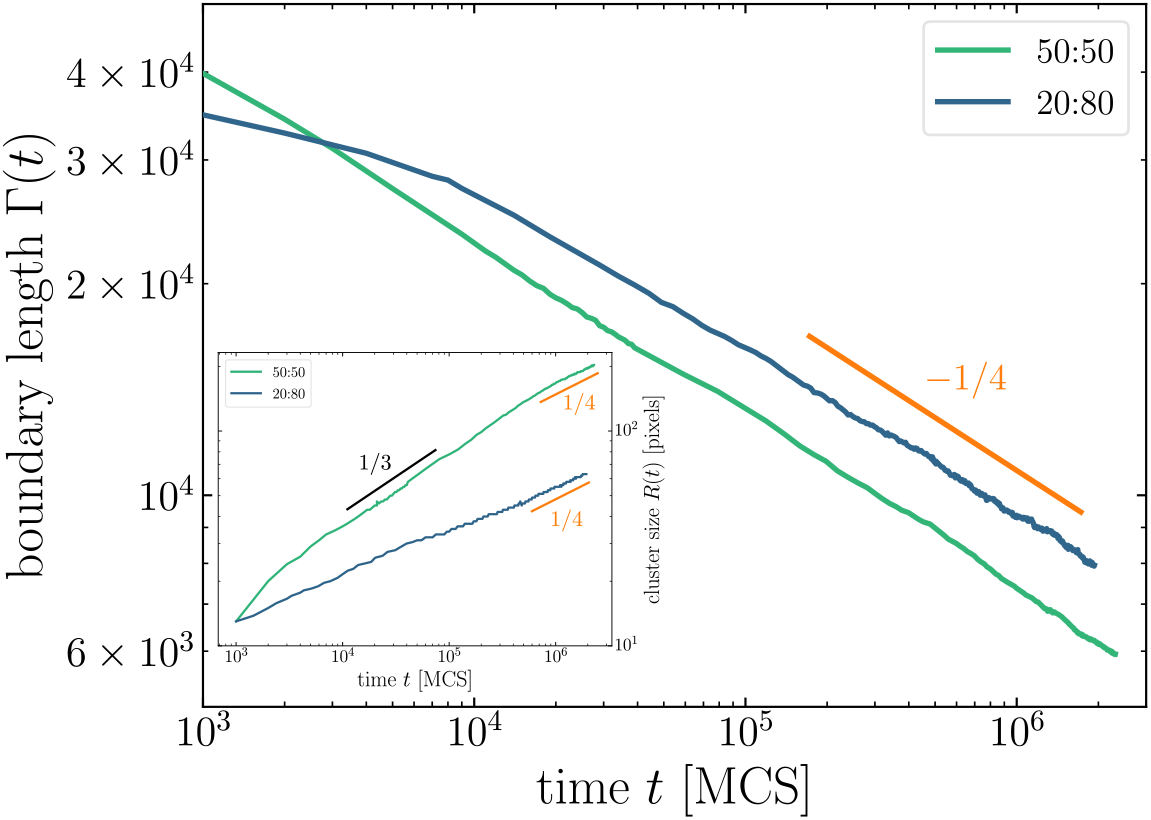
Time evolution of the boundary length Γ(*t*) for even (50:50) and uneven (20:80) mixtures of 4.5×10^4^ endodermal and ectodermal cells (log-log scale). Inset: time evolution of the cluster size *R*(*t*) for the same systems.

We also compare the evolution of the typical *cluster size R*(*t*) for the two mixture ratios. Following [27], we define *R*(*t*) as the position of the first zero of the correlation function

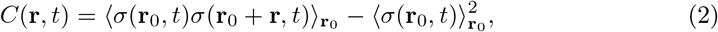

where *σ*(***r***_0_, *t*) is the value at time *t* of the lattice site located at ***r***_0_, and 〈*…*〉**r**_0_ denotes the average over all lattice site locations ***r***_0_. Typical correlation functions are shown in the inset of Fig. 5. In agreement with [9], when they are plotted with the rescaled distance *r/R*(*t*), they all superimpose on a master curve, which shows that the system rescaled by *R*(*t*) exhibits the same statistical property. Note that boundary length and cluster size are related for circular clusters: Γ(*t*) ~ *N* (*t*)*R*(*t*), where *N* (*t*) is the number of clusters, whereas mass conservation implies *N ~ R^−2^*, yielding Γ ~ *R^−1^*. The evolution of *R*(*t*) for the two mixture ratios is shown in the inset of Fig. 4. On long timescales, both curves follow the scaling regime *R*(*t*) ~ *t^m^* with *m* = 1/4, indicating that clusters have circular shapes in this domain. Note however that for the uneven mixture this scaling regime is reached much more rapidly. For the even mixture, the curve follows a quite long transitory regime *R*(*t*) ~ *t*^1/3^, indicating that clusters are not rounded yet.

Our results then suggest a universal scaling *n* = *m* = 1/4 on long timescales, regardless of the mixture ratio, in contradiction with results of Nakajima and Ishihara [9]. Our exponent values also differ from those reported more recently by Cochet-Escartin *et al.* [8]. We discuss in the next section the origins of these discrepancies.

**Fig 5.**
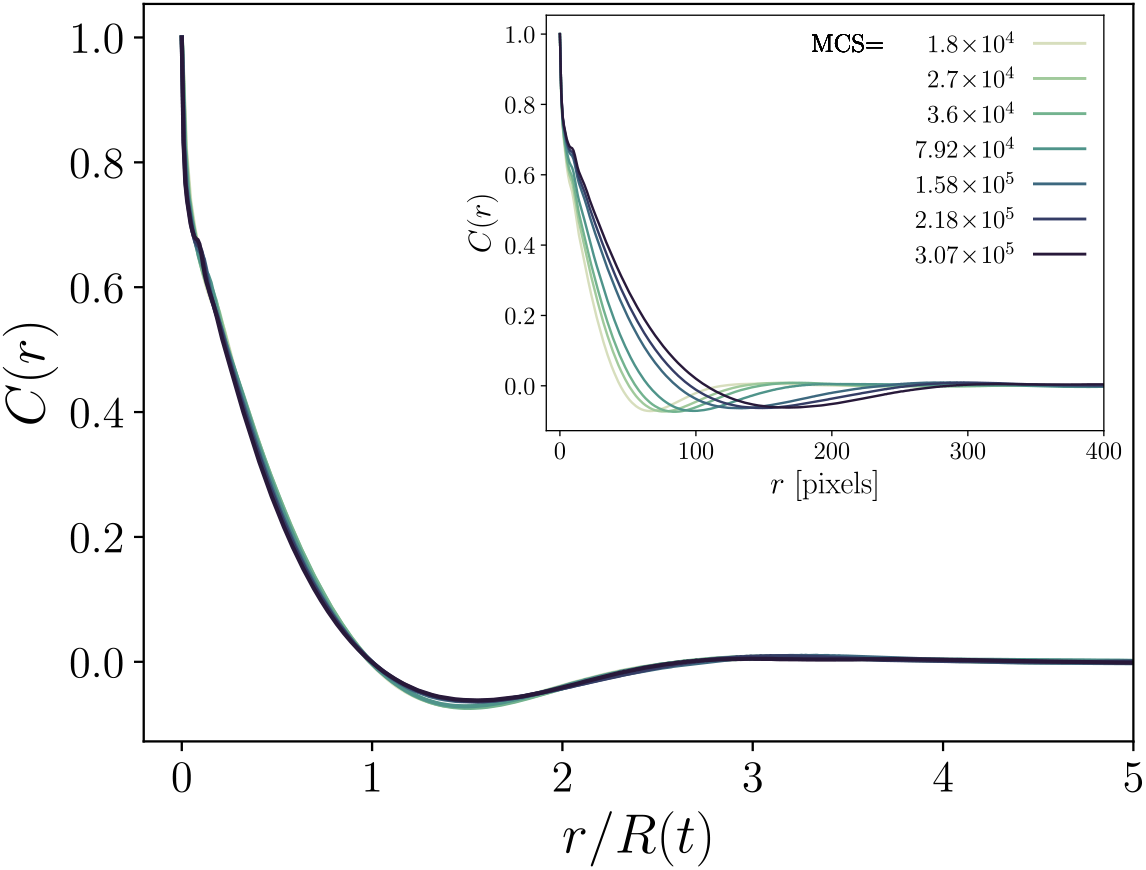
Correlation functions *C*(*r, t*) as functions of the rescaled distance *r/R*(*t*) for several time points, where *R*(*t*) is determined as the position of the first zero of the raw correlation function at time *t*, shown in the inset.

## Discussion

There are two major mechanisms for phase demixing and domain growth. The first one consists in the “evaporation” then “condensation” of the units (atoms, or here, cells) from the smaller clusters to the larger ones. In this *evaporation-and-condensation* mechanism, centers of mass of clusters are fixed. The second mechanism is the diffusion then coalescence of clusters. In this *diffusion-and-coalescence* mechanism, centers of mass of clusters are free to move. In real systems, combination of these two idealized mechanisms can occur. For each of these two mechanisms, kinetics result from two competing processes: the growth rate of the clusters on one side, and the rounding of their interfaces on the other side. The well known Ostwald ripening corresponds to evaporation-and-condensation mechanism when kinetics is limited by the transfer of material from smaller to larger clusters, which then have a rounded shape. Lifshitz, Sliozov and Wagner (LSW) theory then predicts that the domain growth follows a power law ~ *t^m^* with exponent *m* = 1/3 independently of space dimension [28]. Although LSW theory assumes that the volume fraction of one of the components of the mixture is vanishingly small, it has been reported to hold for larger volume fraction as well [28, 29]. As for the diffusion-and-coalescence mechanism, it is well-established [30–32] that in the diffusion-limited regime – in which clusters have rounded shape – domain growth also follows a power-law with exponent *n* = 1/(2 − *dα*), where *d* the space dimension and *α* is the scaling factor between diffusing velocity and mass of aggregates. For cells it is generally accepted that *α* = − 1 [9, 32], yielding *n* = 1/4 at two dimensions.

These results suggest that in our simulations cell sorting is achieved through diffusion-and-coalescence mechanism and that on long times the kinetics is limited by the diffusion of (rounded) clusters, both for even and uneven mixture ratios. Actually, this should come as no surprise, since cluster diffusion has a slower kinetics than cluster rounding, which generally is characterized by an exponential rather than algebraic law [7, 8] – although Nakajima and Ishihara attribute the *n* = 1/3 power law they observed for even mixture to the cluster rounding process [9].

Actually, the *n* = 1/3 power-law reported by Nakajima and Ishihara suggests that the diffusion process is frozen for even mixtures. Presumably, this is a consequence of the periodic conditions they use in their simulations, combined with finite size effects. Indeed, we noticed in our simulations that for even mixtures there is a long transient regime during which cluster growth process is faster than cluster rounding up process, because of the high concentration of clusters. This results in the formation of intertwined structures (which are not all connected in large enough systems), as shown in Fig. (1b). We believe that the *m* = 1/3 power-law variation of the cluster size *R*(*t*) we observe (see inset of Fig. 4) corresponds to this transient regime where clusters grow in size but are not circular. As these clusters keep growing, their diffusion slows down. Eventually, on long times, rounding is faster and the kinetics follows the *n* = 1/4 power-law, typical of the regime of diffusion and coalescence of circular clusters.

When simulations are performed with periodic boundary conditions, these intertwined structures rapidly span the simulation lattice and close on themselves via the periodic boundary conditions, as illustrated in Fig. 6. Such spanning clusters then stop diffusing immediately (as their displacement requires a global move of the whole system), and hinder interface smoothing, because of a competition between spanning clusters that tend to form stripes with flat boundaries, and smaller clusters that tend to form circular boundaries. To test whether the *n* = 1/3 scaling law reported by Nakajima and Ishihara for even mixture reflects the kinetics of cluster boundary smoothing or the transient growth of intertwined structures as we believe, it would be interesting to measure the evolution of the mean cluster area together with the mean radius: during the smoothing process, the mean cluster area should remain constant.

**Fig 6.**
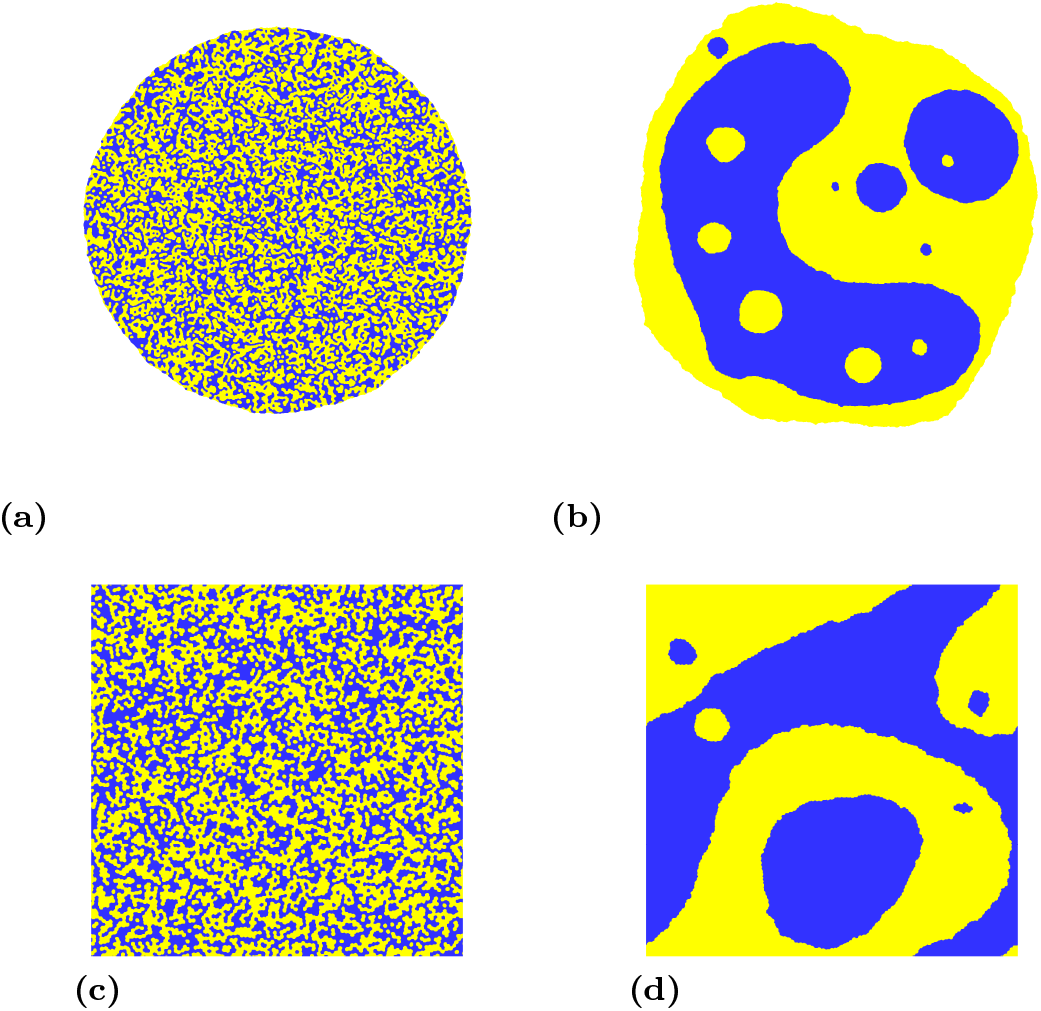
Time evolution of a cluster of 20×10^3^ cells with 50%-50% proportion of endodermal and ectodermal cells: (a)-(b) cluster surrounded by medium (free boundary conditions) at *t* = 10^3^ MCS and *t* = 6×10^6^ MCS; (c)-(d) cluster with periodic boundary conditions at same time points.

In addition to boundary conditions, initial geometry of the aggregate and dimension of space also affect the kinetics of cell sorting, as pointed out by Cochet-Escartin *et al* [8]. In their study, cells were initially placed on a sheet in a 3D space. The exponents they observed in this configuration, either experimentally or numerically, were significantly higher than 1/4. However, the initial flat configuration cannot affect the long time scaling of domain growth, and the evolution in a 3D space should rather decrease the exponent (passing, in the diffusion-limited regime, from *n* = 1/(2 − *dα*) = 1/4 in 2D to *n* = 1/5 in 3D). Moreover, the exponents for Γ(*t*) and *R*(*t*) were quite different in their experimental data, suggesting that clusters did not have time to round up. Finite size effects, but also hydrodynamic couplings (for experiments) [33, 34], could explain the larger exponent values they obtained.

The good agreement between the scaling law reported in the present study and in [9] for uneven binary mixtures confirms that the real time-correspondence of 1 Monte Carlo Step (MCS) for the standard CPM algorithm and ours, which prevents fragmentation, are proportional to each other *τ′*_1MCS_ = *ατ*_1MCS_, where the prefactor *α* depends only on the parameters that characterize the system (here *J_ij_*, *B* and *A*_0_). This linear relationship can be easily explained: as the frequency of fragmentation events that occur with the standard algorithm is homogeneous in time, the amount of time spent in the creation of fragments is homogeneous all along the sorting process.

## Conclusion

Using a modified CPM algorithm that forbids cell fragmentation and increases computational efficiency, we were able to assess the kinetics of cell sorting driven by differential adhesion on very large systems (up to 320, 000 cells). Our results show that on long times, kinetics is controlled by diffusion of rounded clusters binary mixture for cell aggregate surrounded by medium. The growth exponent is *n* = 1/4 for two-dimensional systems, regardless of the mixture ratios of the two cell types. We highlighted the effect of boundary conditions on the scaling law observed on long times, and showed that it explains the apparent contradiction with previous studies.

As a final note, this study confirms that cell sorting driven by differential adhesion has a slow kinetics, and thus is efficient for small multicellular systems only. For larger systems, other mechanisms, such has chimiotaxis, must be involved in the spontaneous segregation of cells.

